# Designing fecal microbiota transplant trials that account for differences in donor stool efficacy

**DOI:** 10.1101/065383

**Authors:** Scott W. Olesen, Thomas Gurry, Eric J. Alm

**Affiliations:** Department of Biological Engineering, Massachusetts Institute of Technology, Cambridge, MA, USA; Center for Microbiome Informatics and Therapeutics, Massachusetts Institute of Technology, Cambridge, MA, USA; The Broad Institute of MIT and Harvard, Cambridge, MA, USA

## Abstract

Fecal microbiota transplantation (FMT) is a highly effective intervention for patients suffering from recurrent *Clostridium difficile*, a common hospital-acquired infection. FMT’s success as a therapy for *C. difficile* has inspired interest in performing clinical trials that experiment with FMT as a therapy for other conditions like inflammatory bowel disease, obesity, diabetes, and Parkinson’s disease. Results from clinical trials that use FMT to treat inflammatory bowel disease suggest that, for at least one condition beyond *C. difficile*, most FMT donors produce stool that is not efficacious. The optimal strategies for identifying and using efficacious donors have not been investigated. We therefore examined the optimal Bayesian response-adaptive strategy for allocating patients to donors and formulated a computationally-tractable myopic heuristic. This heuristic computes the probability that a donor is efficacious by updating prior expectations about the efficacy of FMT, the placebo rate, and the fraction of donors that produce efficacious stool. In simulations designed to mimic a recent FMT clinical trial, for which traditional power calculations predict ~100% statistical power, we found that accounting for differences in donor stool efficacy reduced the predicted statistical power to ~9%. For these simulations, using the heuristic Bayesian allocation strategy more than quadrupled the statistical power to ~39%. We use the results of similar simulations to make recommendations about the number of patients, number of donors, and choice of clinical endpoint that clinical trials should use to optimize their ability to detect if FMT is effective for treating a condition.

## 2 Introduction

Fecal microbiota transplant (FMT), the transfer of stool from a healthy person into an ill person’s gut, is a highly effective treatment for recurrent *Clostridium difficile* infections, which kill 30,000 Americans a year. Despite FMT’s efficacy and increasingly widespread use, the biological mechanism by which FMT cures the infection is not fully understood [1, 2, 3, 4, 5, 6, 7]. FMT’s success in treating *C. difficile* has generated interest in experiments that use FMT to treat other conditions related to the gut and the gut-28 associated microbiota [8, 9]. However, emerging evidence suggests using FMT for these other diseases will be more challenging. Notably, in a recent study by Moayyedi *et al.* [10] that used FMT to treat ulcerative colitis, patients appeared to respond to stool from only one of the six stool donors. Stool from all other donors was no more efficacious than placebo. These results suggest that the effectiveness of FMT can depend strongly on the choice of stool donor.

Ideally, information collected before or during a clinical trial could be used to identify which donors are efficacious. Predictions about a donor’s efficacy could be repeatedly updated depending on that donor’s performance, the performance of other donors, and prior expectations about donor efficacy and heterogeneity. There is, however, not enough clinical information about this variability or biological information about the mechanism by which FMT treats disease to create or validate a model for any particular indication. We therefore present a general model of differences in donor stool efficacy that is not specific to a clinical indication or biomarker. Using this model, we describe the optimal, adaptive, Bayesian strategy for allocating donors and then formulate a computationally-tractable myopic Bayesian allocation heuristic.

Using simulations of clinical trials, we show that differences in donor efficacy and the trial’s strategy for allocating donors can have substantial impacts on the trial’s statistical power. We compare the performance of non-adaptive approaches for allocating patients to donors against a variation of a previously-studied adaptive algorithm (a “play the winner” strategy [11, 12, 13]) and our own Bayesian heuristic. We find that, in many cases, traditional non-adaptive allocation strategies are likely to falsely conclude that FMT in inefficacious. Adaptive strategies, however, can substantially increase a trial’s ability to detect if FMT is efficacious.

## 3 Model of differences in donor stool efficacy

The results of the trial reported in Moayyedi *et al.* [10] raise the possibility that only some donors produce stool that is efficacious for treating ulcerative colitis, type of a inflammatory bowel disease. This possibility—that donors produce stool that is either efficacious or not—is a special case of a more general model of donor-patient interaction.

In this general model, a patient in the treatment arm can respond to the treatment under one of three scenarios:

1. the patient failed to respond to the FMT itself but did respond to the placebo effect,
2. the patient responded to the FMT and would have responded to the placebo effect if they had been given only a placebo, or
3. the patient responded to the FMT and would *not* have responded to a placebo.

We account for these three possibilities by treating the two effects—an effect from stool and the placebo effect—as independent. Specifically, we say that stool has some “active ingredient” that, in the absence of a placebo effect, could cause a patient to respond.

In theory, stool from different donors could have different ingredient efficacies for different patients. Assuming that there are discrete classes of patients and donors, we write the ingredient efficacy of stool from a donor in class *d* for a patient in class *t* as 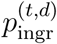. The probability 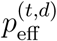 that a patient in class *t* will respond to treatment (i.e., the treatment efficacy) using a donor in class d depends on 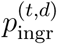 and the probability *p*_pl_ that that patient will respond to the placebo effect:

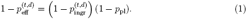

The “matching” between patients and donors, if it exists, is not well understood and cannot yet be predicted. Furthermore, to minimize the patients’ exposure to unknown pathogens, current FMT trials usually treat each patient with stool from only one donor. Thus, each patient is selected randomly from the different patient classes, so the observed treatment efficacy 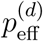 is actually the expected value of 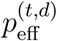 over the different patient classes *t* and patient placebo rates *p*_pl_. Let 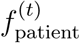 be the fraction of patients that are in class *t* and *f*_pl_ be the probability distribution function of the placebo rates so that:

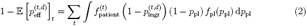

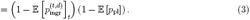

Due to this limitation, we will model only the variables 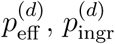 and *p*_pl_ with the understanding that these variables are actually the mean values over distributions of patient characteristics.

The study by Moayyedi *et al*. [10] suggests that the majority of donors produce stool that has no efficacy beyond placebo, that is, that there is a class of donors for which 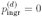. In light of the paucity of the available data about donor qualities, we made a further simplification to the model: we assert that a fraction *f*_eff_ of donors have ingredient efficacy *p*_ingr_ > 0 and the remaining donors have *p*_ingr_ = 0 (i.e., *p*_eff_ = *p*_pl_). This final model is summarized in Figure 1.

**Figure. 1.**
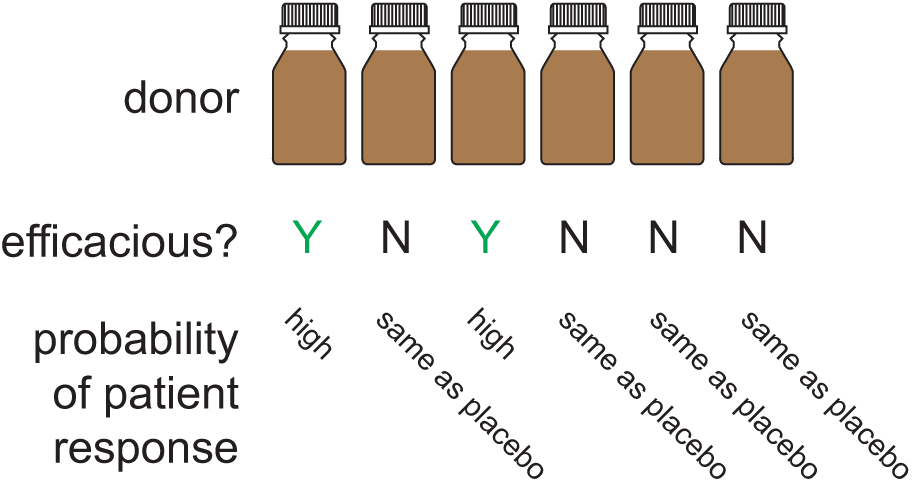
The model of differences in donor efficacy. In the model, donors are efficacious or not. Patients respond to FMT from an efficacious stool donor with probability *p*_eff_. An FMT from an inefficacious stool donor is considered identical to a placebo, i.e., patients respond with probability *p*_pl_. The fraction of donors in the general donor population that are efficacious is *f*_eff_.

### 3.1 Bayesian model formulation

Adaptive donor assignment strategies aim to use the information derived from the patients’ outcomes—and possibly some *a priori* beliefs about the values of the model parameters—to make decisions about how to assign donors. The donor assignment problem, then, is amenable to a Bayesian treatment, which will model the three parameters described above as well as the “qualities” ***q*** ∈ {0, 1}^*D*^ of the *D* donors, where *q_i_* = 0 indicates that donor *i* produces inefficacious (i.e., placebo-equivalent) stool and *q_i_* = 1 indicates that that donor produces efficacious stool.

As described in the Results, we found that the results of simulations using this model were mostly insensitive to the choice of prior distribution on the parameters (*P*_pl_, *P*_ingr_, *f*_eff_), so in most cases we used a simple uniform prior on these values. In one set of simulations, we articulated these priors using beta distributions:

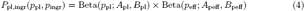

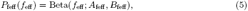

where the hyperparameters *A*_(·)_ and *B*_(·)_ can be drawn from prior clinical trial results. We chose to model the prior on *p*_ingr_ via a beta distribution on *p*_eff_ because *p*_eff_ is directly observable in terms of patient outcomes, while *p*_ingr_ can only be back-computed from *p*_eff_ and *p*_pl_ via equation (1). The clinical trial results for Moayyedi *et al*. [10] and their translation into the model parameters and prior hyperparameters are shown in Table 1.

**Table 1.** Point estimates and hyperparameters for model parameters using clinical data [10]. *A* and *B* are hyperparameters for a beta distribution.

| Clinical trial result | Parameter | Point estimate | $A$ | $B$ |
| --- | --- | --- | --- | --- |
| 2 of 37 in placebo arm responded | $p_{\text{pl}}$ | $2/37 = 0.054$ | 2 | 35 |
| 1 of 6 donors appeared efficacious | $f_{\text{eff}}$ | $1/6 = 0.17$ | 1 | 5 |
| 7 of 18 patients allocated to the efficacious donor responded | $p_{\text{eff}}$ | $7/18 = 0.39$ | 7 | 11 |

We model the prior on the donor qualities ***q*** as a function of *f*_eff_, the fraction of donors in the general population that are efficacious:

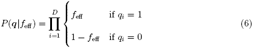

The data used to update the priors are the numbers of patients treated successfully and unsuccessfully for each donor. The likelihood of the data ***X*** given these parameters is

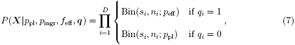

where Bin(*s,n*; *p*) is the binomial probability mass function with *s* successes among *n* total trials with success probability *p, s_i_* is the number of patients successfully treated with stool from donor *i*, and *n_i_* is the total number of patients treated with stool from donor *i*.

Combining equations (6) and (7) shows that the posterior probability of the parameters ***q*** and ***π*** ≡ (*P*_pl_, *P*_ingr_, *f*_eff_) is

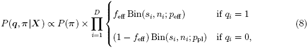

where *P*(***π***) is the prior. The derivation of the posterior predictive probability of a patient responding to treatment with stool from a given donor is in the Appendix.

We found no closed-form solution to the integrals required to compute the predictive posterior probabilities, which we instead evaluated using numerical methods. Specifically, we used Monte Carlo integration with Suave (SUbregion-Adaptive VEgas), an importance sampling method combined with a globally adaptive subdivision strategy. Sampling for this integral was performed with Sobol pseudo-random numbers. The integrator was implemented in C++ using the Cuba package [14]. Wrappers for the integration routine were implemented in Python 3 and simulations were then parallelized to run on multiple cores to optimize computational run time [15].

### 3.2 Optimal donor allocation

Given the priors on the model parameters and the observed patient outcomes, what is the optimal choice for the next donor? This question is similar to other “bandit” problems [16]. Answering it requires evaluating the ramifications that a donor choice will have for the next patient as well as for the trial as a whole.

Let a trial *state* be a trial with some set of observed patient outcomes. Given a donor choice, each trial state has two *child* states: one in which the patient assigned to that donor responded to treatment, one in which the patient did not respond. The optimal donor choice is the one with maximum expected utility *U*, where the utility reflects the outcome of the entire trial. The utility of a donor choice is

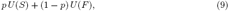

where *p* is the probability of a patient responding to treatment from that donor, *S* is the child state with a success from that donor, and *F* is the child state with a new failed patient.

FMT trials are often designed with a patient horizon (i.e., they will terminate after *N* patients have been treated), and it is natural to associate utilities with the terminal trial states. For example, it would be sensible to take the actions that are most likely to lead to a trial with a significant *p*-value as measured by frequentist statistics, so every terminal node with a significant *p*-value would have utility 1 and the others would have utility 0.

Equation (9) shows that the optimal donor choice at a non-terminal trial state depends only on the utility of its child nodes and each donor’s probability of a successful treatment given that trial state. Thus, the optimal choice at an intermediate trial state can be found by backward induction:

1. Associate each terminal trial state (i.e., those with *N* patients treated) with a utility.
2. For each trial state with *N* − 1 patients treated, compute the expected utility of each donor choice. Associate that state with the expected utility of the donor that maximizes that value.
3. Iterate backwards until the intermediate state has been reached. Select the donor with the maximum expected utility.

An algorithm for counting the number of unique trial states as function of the a number *N* of patients treated, derived in the Appendix, shows that a moderate-size trial with *N* = 30 has more than 10 million terminal trial states. Unfortunately, evaluating the posterior predictive probability that a patient will respond to treatment from a donor 14 9 is computationally intensive, making this optimal approach infeasible.

### 3.3 Deterministic myopic allocation heuristic

In light of the intractability of the optimal solution, we formulated a myopic heuristic that, rather than optimizing the expected utility of the entire trial, simply assigns the next patient to the donor that maximizes the probability that that patient will have a successful outcome. At each step in the trial, the predictive posterior probability of a successful patient outcome is computed for each donor, and the donor with the maximum probability is used. (In the case of a tie, a donor is chosen at random.)

## 4 Methods

### 4.1 Comparison of donor allocation strategies

To evaluate the effectiveness of our myopic heuristic, we performed simulations of clinical trials using four different allocation strategies. The first two strategies, block allocation and random allocation, are non-adaptive. The other two strategies, arandomized urnbased strategy and the myopic Bayesian heuristic, are response-adaptive.

#### Block allocation

In a block allocation, patients are evenly allocated to donors. For example, if there are 30 patients and 6 donors, the first 5 patients are treated with the first donor, the second 5 patients are treated with the second donor, etc. (In clinical practice, patients would be randomized within the blocks.)

#### Random allocation

In a random allocation, patients are allocated to donors at random. (On average, random allocations are similar to block allocations, so these two types of non-adaptive simulation yield similar results.)

#### Urn-based allocation

Our urn-based allocation strategy is a variation on the “play-the-winner” strategies designed and studied as an ethical [17] and statistically-rigorous way to decide how to allocate patients to a treatment arm when a trial includes more than one treatment arm [18, 19, 20]. In this study, we used the generalized Póolya’s urn [21] with parameters *w* = 1, *α* = 3, *β* = 0, and without replacing the drawn ball.

#### Myopic Bayesian heuristic

This is the strategy described in the previous section. Except when noted, a uniform prior on the three parameters (*P*_pl_, *P*_ingr_, *f*_eff_) was used.

## 4.2 Simulated clinical trials

The expected fraction of patients allocated to efficacious donors and the statistical power of clinical trials using different donor allocation strategies were estimated using simulated clinical trials. In each simulation, the three model parameters (*P*_pl_, *P*_eff_, *f*_eff_) and the number of patients in the trial were fixed. For each combination of the trial parameters, 10,000 lists of 6 donors each were randomly generated. Donors were designated as efficacious or not efficacious by random chance according to the frequency of efficacious donors *f*_eff_. The same donor lists were used for simulations for each of the allocation strategies.

In one set of simulations, the number of donors was varied among 1, 3, 5, 10, 15, and 30. For those simulations, lists of 30 donors were generated for each parameter set and trial iteration. The lists were truncated for the simulations using less than 30 donors.

For each allocation strategy and donor list, atrial was simulated. In each simulation, apatient allocated to an efficacious donor responds to the treatment with probability *P*_eff_. Patients allocated to inefficacious donors or to the placebo arm respond with probability *P*_pl_. For adaptive allocations, the outcomes from all the previous patients treatment were determined before the donor for the next patient was selected. An equal number of patients was allocated to the treatment and placebo arms. (This equal allocation to the control arm was intended to preserve statistical power, which would probably have decreased if the myopic algorithm were allowed to allocate patients away from the placebo arm [22, 23].)

### Clinically-relevant parameter values

Simulations were performed for all combinations of parameter values selected to reflect clinically-relevant possibilities:

- The placebo rate *P*_pl_ is either 0.05 (a low placebo rate consistent with stringent, objective outcomes; e.g., endoscopic Mayo score [10, 24]) or 0.25 (ahigh placebo rate consistent with self-reported, subjective outcomes [25, 26]).
- The efficacy *P*_eff_ of efficacious donors is either 0.4 (similar to the value in Table 1) or 0.95 (efficacy of FMT to treat *C. difficile* infection).
- The frequency *f*_eff_ of efficacious donors is either 0.15 (similar to the value in Table 1) or 0.9 (reflective of the fact that almost any well-screened donor produces stool that can successfully treat *C. difficile* infection).
- The number of patients in each of the treatment and control arms is 15, 30, or 60, corresponding to a range of patient numbers typical for Phase I and small Phase II clinical trials.

Of these combinations, the set of values most similar to the one in Table 1 is *P*_pl_ = 0.05, *P*_eff_ = 0.4, *f*_eff_ = 0.15, *N*_patients_ = 30.

### Computing statistical power

After determining the outcome of all the patients in the trial, the *p*-value of a one-sided Fisher’s exact test (asserting that the response rate in the treatment arm was greater) was calculated. The proportion of simulations that produced *p* < 0.05 was the estimate of the statistical power for that allocation strategy under those trial parameters. Confidence intervals were calculated using the method of Clopper and Pearson [27]. Values are rounded to two or three significant digits.

## 5 Results

### 5.1 Trials using adaptive strategies allocate more patients to efficacious donors

The purpose of adaptive donor allocation strategies is to identify and use efficacious 223 donors. We therefore expected that simulated trials that use adaptive strategies would allocate more patients to efficacious donors (compared to simulated trials that used the block or random donor allocation strategies).

For every parameter set simulated, the average fraction of patients allocated to efficacious donors was greater in the adaptive strategies (urn-based and myopic Bayesian) than in the non-adaptive strategies (block and random; Table S1). The two non-adaptive allocation strategies performed almost identically: for each parameter set, their results differed by less than 1 percentage point. The two adaptive strategies performed similarly: for half of the parameter sets, their results differed by less than 2 percentage points. In the remaining parameter sets, their results varied by between 2 and 9 percentage points.

When efficacious donors are common (*f*_eff_ = 0.9), the adaptive and non-adaptive strategies performed similarly (Fig. 2). In other cases, the performance of the two strategies differed substantially. For example, for the parameterization most similar to the one in Table 1, the random strategy allocated 15% of patients to efficacious donors while the myopic Bayesian strategy allocated 41% of patients to efficacious donors.

**Figure. 2.**
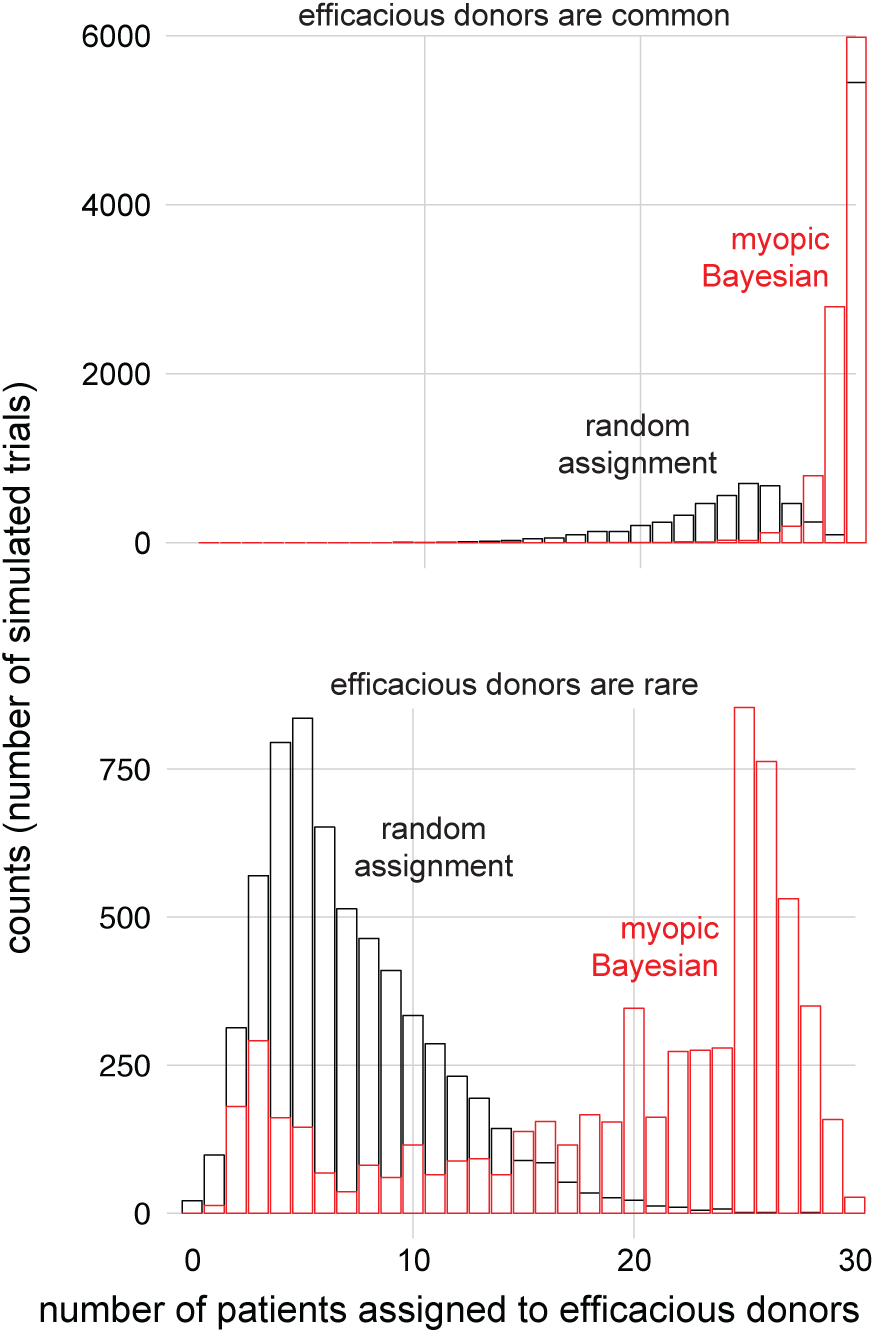
The Bayesian myopic strategy allocates more patients to efficacious donors. (top) For the parameterization most similar to the one in Table 1 (top), the Bayesian myopic heuristic (red) allocated more patients to efficacious donors than random allocation (black) did. When efficacious donors are common (bottom; same parameters as top but with *f*_eff_ = 0.9), the two strategies allocate similar numbers of patients to efficacious donors. For visual clarity, only trials in which there was at least one efficacious donor are shown.

### 5.2 Trials using adaptive allocation have higher statistical power

Because trials that used the adaptive donor allocation strategies allocated more patients to efficacious donors than the trials that used the non-adaptive strategies, we expected that trials using adaptive strategies would have greater statistical power.

The adaptive strategies consistently yielded higher statistical powers than the nonadaptive strategies (Table 2). When efficacious donors are rare, the performance gap is larger. For example, for the parameterization most similar to the one in Table 1, a trial that uses random allocation is expected to have 9% power, while the myopic Bayesian strategy can deliver 39% power. The gap in performance is smallest when selecting a donor at random is likely to yield an efficacious donor: among the trials with *f*_eff_ = 0.9, the adaptive and non-adaptive statistical powers differed by less than 6 percentage points.

Low statistical powers when *f*_eff_ is small are likely due to the fact that if *all* the available donors are not efficacious, then no allocation strategy should make a trial achieve significance. For example, if only 15% of donors are efficacious (*f*_eff_ = 0.15) and there 252 are only 6 donors (the number used in these simulations), then we expect that 38% of trials will have no good donors (using the binomial distribution function). We therefore separately analyzed the simulated trials in which no donors were efficacious and the simulated trials in which at least one donor was efficacious (Table S2). When no donors are efficacious, trials with adaptive or non-adaptive strategies have ~0% power. Among trials with at least one efficacious donor, the difference in statistical power between adaptive and non-adaptive strategies is greater than the difference computed using the results of all trials.

**Table 2.**
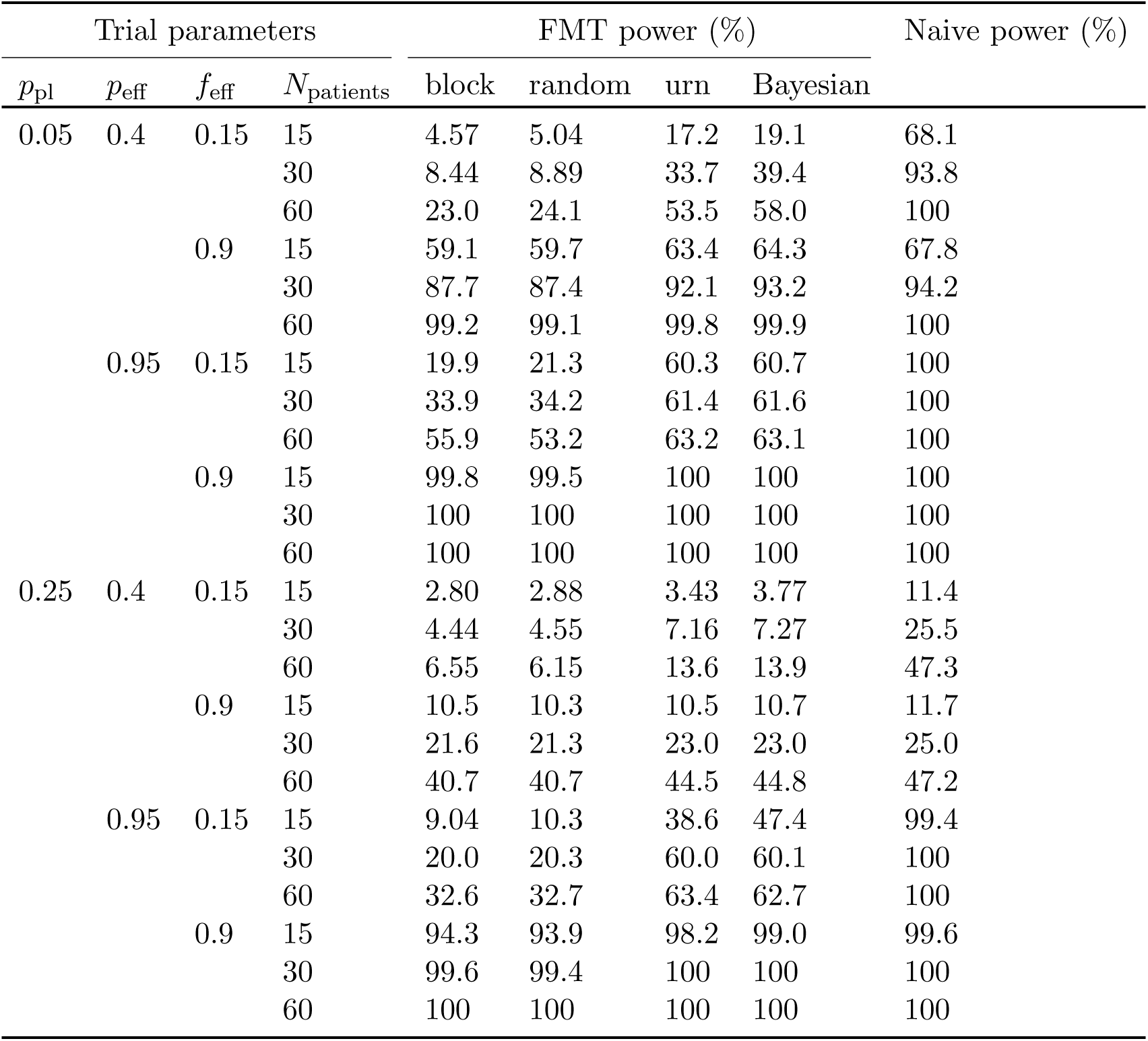
Adaptive strategies yield clinical trials with higher statistical power. “FMT power” is the power computed by simulating the results of trials that would occur if the frequency of efficacious donors if *f*_eff_. “Naive power” is the power computed in the situation in which all donors are efficacious (i.e., *f*_eff_ = 1.0). All 95% confidence intervals on these values are within 1 percentage point of the reported value and are not shown.

Conversely, the power computed in traditional calculations that do not account for differences in donor efficacy (i.e., that assume that all donors are efficacious, or equivalently *f*_eff_ =1.0) is, in many cases, substantially higher than the power computed when accounting for differences in donor efficacy (Table 2, column “Naive powers”). For example, for the parameter set most similar to the one in Table 1, the naive calculation predicts 94% power, but the calculation that accounts for differences in donor efficacy predicts only 9% power for non-adaptive allocation strategies. The differences between the powers computed by the naive method and our approach is largest when *f*_eff_ is small.

### 5.3 Performance of adaptive strategies depend on their parameterization

In these simulations, we varied the actual values of *P*_pl_ and *P*_eff_ but we always initialized the adaptive algorithms the same ways. To determine the sensitivity of the adaptive allocation algorithms’ performance to their initialization, we simulated trials in which the actual model parameters were fixed but the algorithms’ initializations varied (Table S3 and Table S4). The myopic Bayesian algorithm’s performance was mostly robust to the parameterization of its prior distribution except when the prior was strong and inaccurate. Accurate priors, weak priors, and uniform priors provide comparable performance. In contrast, the urn algorithm delivered widely varying powers, from 15% to 40%, depending on its parameterization.

### 5.4 Increasing the number of available donors benefits the adaptive Bayesian strategy

Increasing the number of available donors increases the probability that at least one of them will be efficacious. We therefore determined, for each donor allocation strategy, the number of available donors that optimized the trial’s expected power. Simulations showed that increasing the size of the donor “pool” almost always increased the power of trials using the myopic Bayesian donor allocation but, depending on the parameter set, could increase or decrease the power of trials using other allocation strategies (Table S5).

Table 3 shows how donor selection strategy and model parameter values affects the optimal number of donors. Notably, when efficacious donors are uncommon (*f*_eff_ = 0.15) and only moderately efficacious (*P*_eff_ = 0.4), the non-adaptive strategies perform opti-290 mally when only one donor is used. In other words, when using non-adaptive strategies in this parameter regime, it is wiser to take the 15% chance of picking a single efficacious donor than it is to distribute patients across many donors, allotting around 15% of them to efficacious donors. In contrast, the myopic Bayesian donor allocation almost always benefits from a larger donor pool.

**Table 3.** The optimal number of donors varies by donor selection strategy and model parameter values. For each parameter value, trials were simulated using 1, 3, 5, 10, 15, and 30 donors. (The number of patients was fixed at 30.) The number of donors that optimized the expected power was identified. If multiple numbers of donors yielded powers within 0.05 of the optimal value, all those numbers are reported as a range.

| Trial parameters | | | Optimal $N_{\text{donors}}$ | | | |
| --- | --- | --- | --- | --- | --- | --- |
| $f_{\text{eff}}$ | $p_{\text{eff}}$ | $p_{\text{pl}}$ | random | block | urn | Bayesian |
| 0.15 | 0.4 | 0.05 | 1 | 1 | 10 | 10–30 |
| 0.15 | 0.4 | 0.25 | 1 | 1 | 5 | 10 |
| 0.15 | 0.95 | 0.05 | 3–15 | 3–5 | 15–30 | 15–30 |
| 0.15 | 0.95 | 0.25 | 3 | 3 | 10 | 10–30 |
| 0.9 | 0.4 | 0.05 | 1–30 | 3–30 | 3–30 | 3–30 |
| 0.9 | 0.4 | 0.25 | 1–30 | 1–30 | 1–30 | 5–30 |
| 0.9 | 0.95 | 0.05 | 3–30 | 3–30 | 3–30 | 3–30 |
| 0.9 | 0.95 | 0.25 | 3–30 | 3–30 | 3–30 | 3–30 |

## 6 Discussion

### 6.1 Model limitations

We chose to use a simple model for a simple use case because, in the absence of data about the treatment histories of hundreds of patients using dozens of donors, we do not believe that more complicated models will be more useful to aiding trial design. The model’s greatest weakness is that it cannot be validated, but it is exactly the model’s purpose to improve the probability of collecting the kind of information that could validate or invalidate it. In light of the dearth of data, we developed a simple model, and it could be that the simplifications we made limit the model’s validity. For example, the model assumes that each donor produces efficacious stool or inefficacious stool (when in fact there is probably day-to-day and donor-to-donor variation in stool efficacy) and that all patients receive one course of treatment (while, say, patients who do not respond to a first treatment might be treated with stool from a different donor).

In our formulation of the model, we only considered optimizing the assignment of patients to donors within the treatment arm, thus “protecting” the size of the placebo arm. Future work could determine if there is a benefit to adaptively assigning patients to the control or treatment arms based on estimations of the donors’ quality. We also assumed in our simulations that the outcome from all the previous patients treatments are known before the donor for the next patient is selected. In reality, patients in an FMT trial overlap. The urn-based method can still be used for overlapping patients [21], but the myopic Bayesian method would require some modification. The urn-based approach is also randomized, which is desirable in clinical trials because it can reduce certain kinds of bias [11]. Future work could adapt the myopic Bayesian heuristic, which is deterministic, for adaptive randomization [28] and for delays between patient allocation and outcome observation.

### 6.2 Differences in donor efficacy should be accounted for in trial design

Our results entail recommendations to clinicians. First, the powers we computed here are, in many cases, well below the powers computed assuming that all donors are efficacious. We therefore encourage researchers to consult our predictions about statistical power 325 when deciding on the size of their trials.

Second, a high placebo rate can substantially decrease the statistical power of an FMT trial. We therefore encourage researchers to use the most stringent outcome measurement possible (e.g., an endoscopic Mayo score for inflammatory bowel disease).

Third, adaptive donor allocation strategies consistently delivered higher statistical power than traditional, non-adaptive approaches. We therefore recommend that researchers use such an adaptive strategy. The urn-based strategy has the advantages that similar response-adaptive strategies may be familiar to clinicians, it is randomized, and it is simple to implement. However, an urn-based strategy needs to be carefully parameterized: a badly-parameterized urn-based strategy performs similarly to random allocation. The adaptive Bayesian donor allocation algorithm performs well even when using the “default” settings (auniform prior) but is complex and deterministic. To fully leverage this strategy, aclinician would need to consult the algorithm’s output after every patient outcome and follow the algorithm’s deterministic instructions, which might introduce bias.

Fourth, adaptive algorithms benefit from having access to a “bank” of 10 or more donors. Researchers hoping to achieve the full benefits of adaptive donor selection must be prepared to change donors multiple times during the trial.

Finally, researchers reporting about FMT trials should include information about the donors, notably how many donors were used and what proportion of patients allotted to each donor responded to treatment. This information will help future researchers account for differences in donor efficacy.

### 6.3 Future research may identify mechanistic explanations of FMT’s efficacy

The adaptive allocation strategies we described here have a narrow aim: to increase the number of successful patient outcomes in a trial. In theory, an adaptive trial design is capable of more. For example, if it were hypothesized that FMT succeeded or failed because of the presence or absence of some particular microbial species in the donor’s stool, then an adaptive trial design could recommend donor choices that aim to identify that critical species.

We did not pursue a hypothesis-centric approach because we believe it is premature. Even the mechanism by which FMT treats *C. difficile*, the most well-studied case, remains unclear. We expect that strong hypotheses about mechanism will come from retroactive comparison of efficacious vs. inefficacious stool *after* clinical trials have definitively show that FMT is effective for treating some disease. Our study aims to do exactly this. Until then, we hope that our results about adaptive donor allocation help more patients benefit from FMT and will help clinicians identify those conditions that FMT can treat.

## 7 Acknowledgements

This material is based on work supported by the National Science Foundation Graduate Research Fellowship under Grant No. 1122374 and by the Center for Microbiome Informatics and Therapeutics at MIT.

## A Mathematical appendix

### A.1 Posterior probability of patient response

Let *S_i_* be the event that the next patient responds to treatment with stool from donor *i*. The posterior predictive probability of *S_i_* is

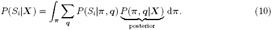

The probability *P*(*S_i_|**π, q***) is simply *P*_eff_ is *q_i_* = 1 and *P*_pl_ if *q_i_* = 0. Thus,

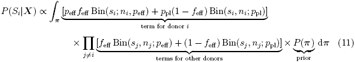

The constant of proportionality is the same for all *i*: the quantity in equation (11) should be divided by

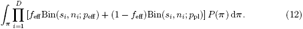

Because the binomials in the equations (11) and (12) have the same constants, Bin(*s, n*; *p*) can be replaced by *p^s^*(1 − *p*)^*f*^ in these equations.

### A.2 Counting trial states

How many unique trial states are there for a given number of patients *N*?

Naively, a trial state can be specified by a string that specifies with donor was used and what outcome was observed (e.g., *AsAfBs* means that donor *A* has a success, then a failure, then donor *B* had a success). A trial with *N* patients can have up to *N* donors, meaning that there are *N^N^* possible chains of donors and 2*^N^* possible chains of outcomes, for (2*N*)*^N^* possible trial states.

However, many of those states are identical. In particular, we consider donors identical unless they have a different number of successful or failed patient outcomes (e.g., *AsAs* is identical to *BsBs*), and we consider trial states identical if the same results are achieved in a different order (e.g., *AsAf* is identical to *AfAs*). Thus, a unique trial set is a set of ordered triples (*d_i_,s_i_,f_i_*), where *d_i_* is the number of donors that each have had *s_i_* successful and *f_i_* failed patient outcomes.

**Theorem 1.** *Let w*(*N, M*) *be the number of trial states with N* ≥ 0 *total patients with at most M* ≥ 0 *patients in any donor*. *W*(*N, M*) *is defined by the recursion*:

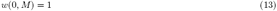

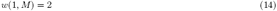

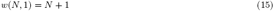

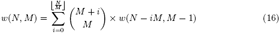

*The number of trial states with N patients is w*(*N, N*).

*Proof*. We then justify each of the recursion rules. For *N* = 0 (i.e., no patients), there is only one trial state: {(0, 0, 0)}. For *N* = 1 (i.e., one patient), there are two trial states: {(1, 1, 0)} and {(1, 0, 1)}. For *M* = 1(i.e., at most one patient per donor), there are *N* + 1 trial states: {(*i*, 1, 0), (*N* − *i*, 0, 1)} for 0 ≤ *i* ≤ *N*. In the general case, let i be the number of donors with *M* patients. This number can range from 0 to 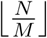. For each of those donors, there are *M* + 1 possible success/failure pairs: (*j, M* − *j*) for 0 ≤ *j* ≤ *M*. Thus, there are 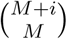 possible trial states for just those *i* donors. This leaves *N* − *iM* patients to be distributed among the remaining donors, who can each have at most *M* −1 patients.

Some representative results are shown in Table S6.

### B Supplementary Tables

**Table S1.** Fraction of patients allocated to efficacious donors. The same simulated trials were analyzed to create Table 2 and this table.

| $p_{\text{pl}}$ | $p_{\text{eff}}$ | $f_{\text{eff}}$ | $N_{\text{patients}}$ | block | | random | | urn | | Bayesian | |
| --- | --- | --- | --- | --- | --- | --- | --- | --- | --- | --- | --- |
|  |  |  |  | mean | s.d. | mean | s.d. | mean | s.d. | mean | s.d. |
| 0.05 | 0.4 | 0.15 | 15 | 0.15 | 0.15 | 0.15 | 0.17 | 0.29 | 0.30 | 0.32 | 0.33 |
|  |  |  | 30 | 0.15 | 0.14 | 0.15 | 0.16 | 0.36 | 0.35 | 0.41 | 0.38 |
|  |  |  | 60 | 0.15 | 0.14 | 0.15 | 0.15 | 0.44 | 0.39 | 0.49 | 0.42 |
|  |  | 0.9 | 15 | 0.9 | 0.12 | 0.9 | 0.14 | 0.95 | 0.075 | 0.96 | 0.068 |
|  |  |  | 30 | 0.9 | 0.12 | 0.9 | 0.13 | 0.97 | 0.051 | 0.98 | 0.041 |
|  |  |  | 60 | 0.9 | 0.12 | 0.9 | 0.13 | 0.98 | 0.03 | 0.99 | 0.022 |
|  | 0.95 | 0.15 | 15 | 0.15 | 0.15 | 0.15 | 0.17 | 0.44 | 0.36 | 0.51 | 0.42 |
|  |  |  | 30 | 0.15 | 0.15 | 0.15 | 0.16 | 0.52 | 0.42 | 0.56 | 0.45 |
|  |  |  | 60 | 0.15 | 0.15 | 0.15 | 0.15 | 0.57 | 0.45 | 0.6 | 0.47 |
|  |  | 0.9 | 15 | 0.9 | 0.13 | 0.9 | 0.14 | 0.97 | 0.047 | 0.99 | 0.028 |
|  |  |  | 30 | 0.9 | 0.12 | 0.9 | 0.13 | 0.98 | 0.027 | 1.0 | 0.013 |
|  |  |  | 60 | 0.9 | 0.12 | 0.9 | 0.13 | 0.99 | 0.014 | 1.0 | 0.0069 |
| 0.25 | 0.4 | 0.15 | 15 | 0.15 | 0.15 | 0.15 | 0.17 | 0.20 | 0.25 | 0.21 | 0.29 |
|  |  |  | 30 | 0.15 | 0.15 | 0.15 | 0.16 | 0.24 | 0.30 | 0.24 | 0.34 |
|  |  |  | 60 | 0.15 | 0.15 | 0.15 | 0.15 | 0.29 | 0.34 | 0.28 | 0.38 |
|  |  | 0.9 | 15 | 0.9 | 0.13 | 0.9 | 0.14 | 0.93 | 0.14 | 0.94 | 0.15 |
|  |  |  | 30 | 0.9 | 0.12 | 0.9 | 0.13 | 0.94 | 0.13 | 0.95 | 0.14 |
|  |  |  | 60 | 0.9 | 0.12 | 0.9 | 0.13 | 0.96 | 0.11 | 0.96 | 0.14 |
|  | 0.95 | 0.15 | 15 | 0.15 | 0.15 | 0.15 | 0.17 | 0.36 | 0.34 | 0.45 | 0.42 |
|  |  |  | 30 | 0.15 | 0.15 | 0.15 | 0.16 | 0.45 | 0.38 | 0.52 | 0.44 |
|  |  |  | 60 | 0.15 | 0.15 | 0.15 | 0.15 | 0.53 | 0.42 | 0.57 | 0.46 |
|  |  | 0.9 | 15 | 0.9 | 0.13 | 0.9 | 0.14 | 0.96 | 0.076 | 0.99 | 0.056 |
|  |  |  | 30 | 0.9 | 0.12 | 0.9 | 0.13 | 0.98 | 0.048 | 0.99 | 0.029 |
|  |  |  | 60 | 0.9 | 0.12 | 0.9 | 0.13 | 0.99 | 0.029 | 1.0 | 0.016 |

**Table S2.**
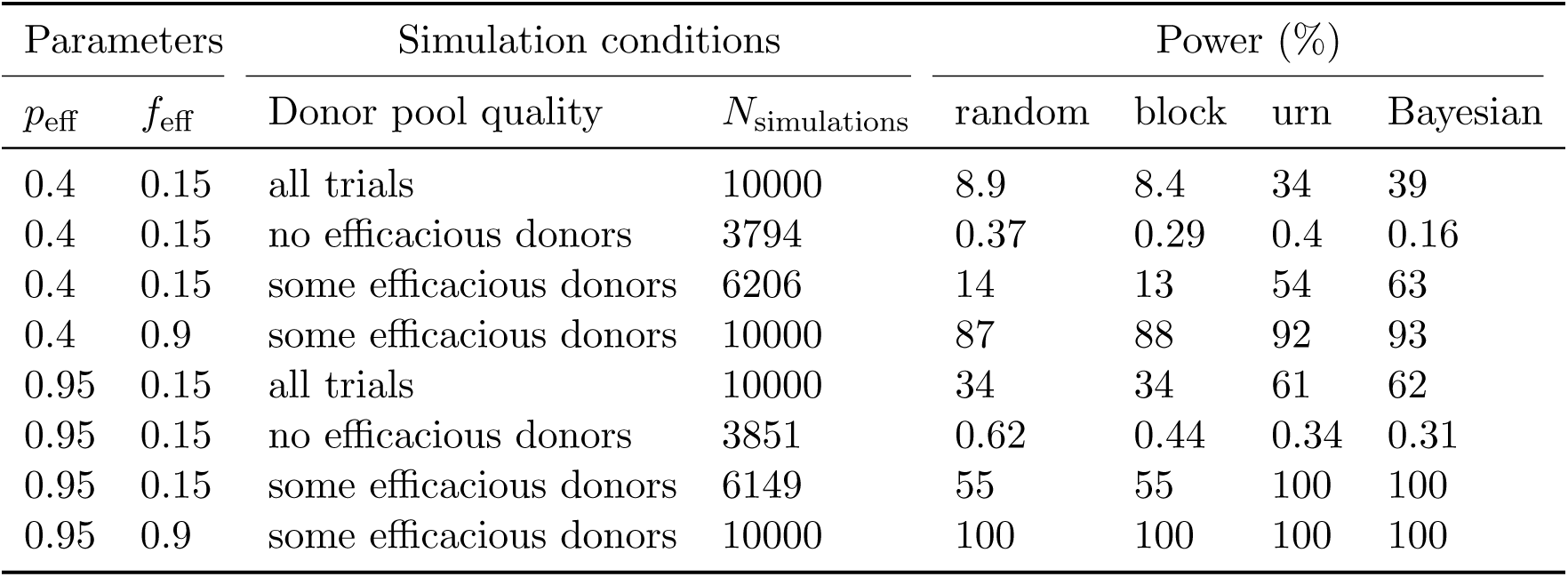
Power of simulated clinical trials, conditioned on presence of efficacious donors. A subset of the data used to create Table 2 (only those simulations with *N*_patients_ = 30 and *p*_pl_ = 0:05) was analyzed by separately estimating the power for the trials in which no donors were efficacious (“no efficacious donors”) and in which at least one donor was efficacious (“some efficacious donors”). “All simulations” shows the same data as in Table 2. For *f*_eff_ = 0:9, none of the 10,000 simulations had a donor pool with no efficacious donors. All 95% confidence intervals are within 1 percentage point of the reported value and are not shown.

| Parameters |  | Simulation conditions |  | Power (%) |  |  |  |
| --- | --- | --- | --- | --- | --- | --- | --- |
| $p_{\text{eff}}$ | $f_{\text{eff}}$ | Donor pool quality | $N_{\text{simulations}}$ | random | block | urn | Bayesian |
| 0.4 | 0.15 | all trials | 10000 | 8.9 | 8.4 | 34 | 39 |
| 0.4 | 0.15 | no efficacious donors | 3794 | 0.37 | 0.29 | 0.4 | 0.16 |
| 0.4 | 0.15 | some efficacious donors | 6206 | 14 | 13 | 54 | 63 |
| 0.4 | 0.9 | some efficacious donors | 10000 | 87 | 88 | 92 | 93 |
| 0.95 | 0.15 | all trials | 10000 | 34 | 34 | 61 | 62 |
| 0.95 | 0.15 | no efficacious donors | 3851 | 0.62 | 0.44 | 0.34 | 0.31 |
| 0.95 | 0.15 | some efficacious donors | 6149 | 55 | 55 | 100 | 100 |
| 0.95 | 0.9 | some efficacious donors | 10000 | 100 | 100 | 100 | 100 |

**Table S3.** Sensitivity of simulated clinical trial power to parameterization of the myopic Bayesian algorithm parameters. Starting from the parameter described in Table 1, 10,000 trials using the myopic Bayesian donor allocation were simulated for each of six different parameterizations of the Bayesian algorithm. “Accurate” means that the prior is centered around the true value (i.e., that 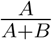 equals the true value); “inaccurate” means that the prior is centered at approximately double the true value (but that *A* + *B* has been held constant). “Strong” means that every hyperparameter is ten-fold greater; “weak” means that every hyperparameter is ten-fold smaller. “Uniform” means a uniform prior was used for all parameters. All 95% confidence intervals are within 1 percentage point of the reported value and are not shown.

| Hyperparameterization | Hyperparameter values |  |  |  |  |  | Power (%) |
| --- | --- | --- | --- | --- | --- | --- | --- |
| | $A_{\text{placebo}}$ | $B_{\text{placebo}}$ | $A_{\text{peff}}$ | $B_{\text{peff}}$ | $A_{\text{feff}}$ | $B_{\text{feff}}$ | |
| From Table 1 | 2 | 35 | 7 | 11 | 1 | 5 | 41.1 |
| Weak, accurate | 0.2 | 3.5 | 0.7 | 1.1 | 0.1 | 0.5 | 41.2 |
| Strong, accurate | 20 | 350 | 70 | 110 | 10 | 50 | 40.4 |
| Weak, inaccurate | 0.4 | 3.3 | 1.4 | 0.4 | 0.2 | 0.4 | 40.9 |
| Strong, inaccurate | 40 | 330 | 140 | 40 | 20 | 40 | 34.4 |
| Uniform | n.a. | n.a. | n.a. | n.a. | n.a. | n.a. | 39.6 |

**Table S4.**
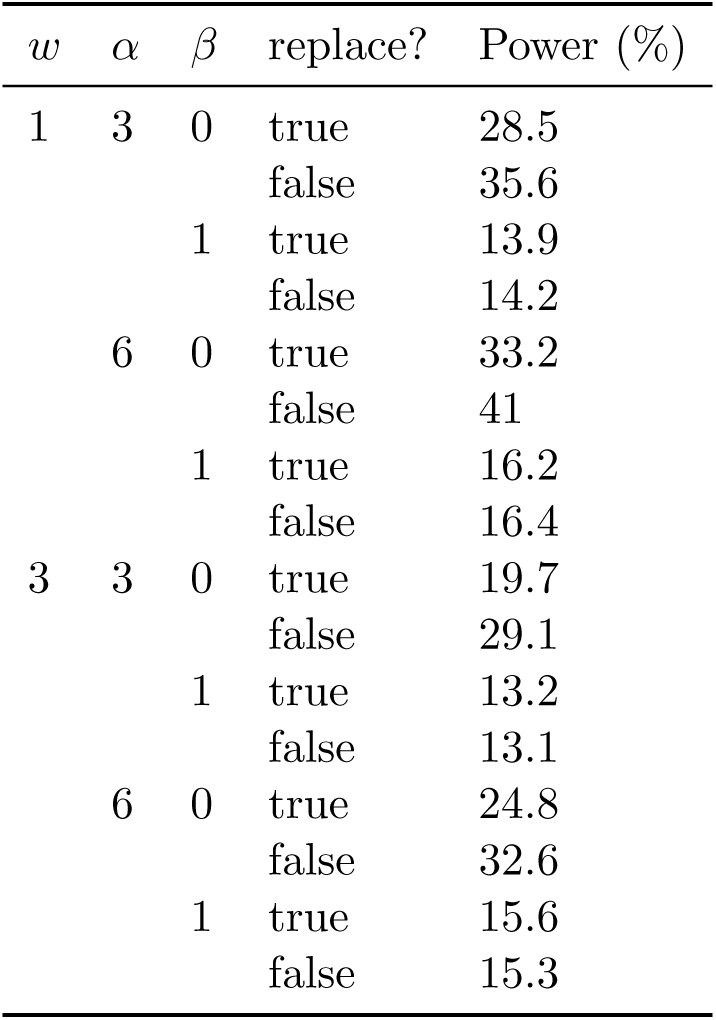
Sensitivity of simulated clinical trial power to urn parameterization. Using the parameter set described in Table 1, 10,000 trials using the urn-based donor allocation were simulated for each of several combinations of the parameters for the urn model as described in [21]. We also vary whether drawn balls are replaced or not. All 95% confidence intervals are within 1 percentage point of the reported value and are not shown.

**Table S5.**
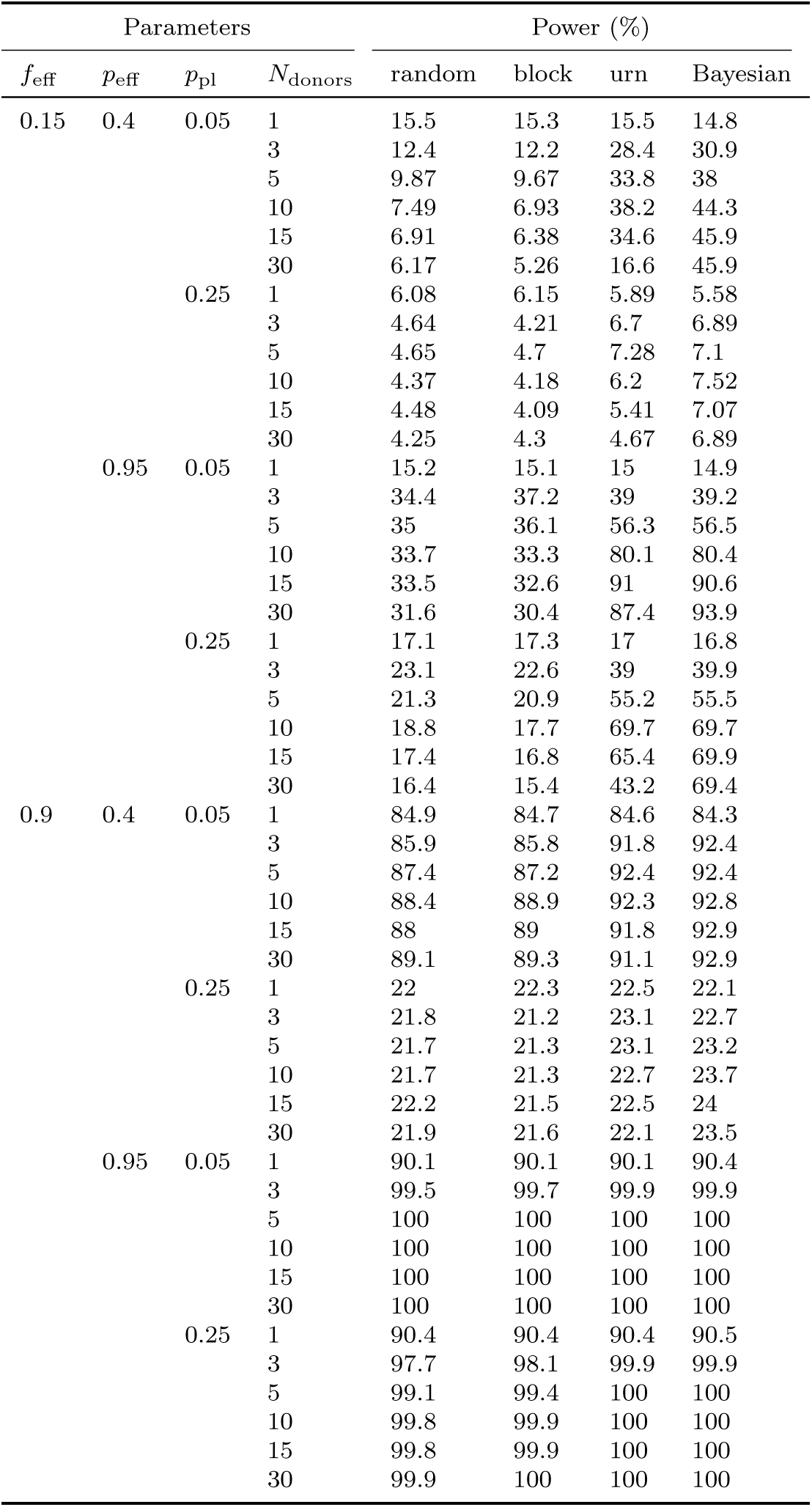
Dependence of simulated clinical trial power on number of available donors. For each parameter value, trials were simulated using 1, 3, 5, 10, 15, and 30 donors. The number of patients was fixed at 30. All 95% confidence intervals are within 1 percentage point of the reported value and are not shown.

**Table S6.** The number of possible trial states for a given number of patients, assuming that the number of available donors is equal to the number of patients.

| # patients | # trial states |
| --- | --- |
| 1 | 2 |
| 2 | 6 |
| 3 | 14 |
| 4 | 33 |
| 5 | 70 |
| 6 | 149 |
| 7 | 298 |
| 8 | 591 |
| 9 | 1,132 |
| 10 | 2,139 |
| 15 | 39,894 |
| 20 | $5.58 \times 10^5$ |
| 25 | $6.39 \times 10^6$ |
| 30 | $6.29 \times 10^7$ |
| 40 | $4.34 \times 10^9$ |
| 50 | $2.14 \times 10^{11}$ |
| 60 | $8.22 \times 10^{12}$ |

## References

[1] Stuart H. Cohen, Dale N. Gerding, Stuart Johnson, Ciarán P. Kelly, Vivian G. Loo, L. Clifford McDonald, Jacques Pepin, and Mark H. Wilcox. Clinical practice guidelines for *Clostridium difficile* infection in adults. Infect Control Hosp Epidemiol, 31(5):431–455, 2010.

[2] Els van Nood, Anne Vrieze, Max Nieuwdorp, Susana Fuentes, Erwin G. Zoetendal, Willem M. de Vos, Caroline E. Visser, Ed J. Kuijper, Joep F. W. M. Bartelsman, Jan G. P. Tijssen, Peter Speelman, Marcel G. W. Dijkgraaf, and Josbert J. Keller. Duodenal infusion of donor feces for recurrent *Clostridium difficile*. N Engl J Med, 368(5):407–415, 2013.

[3] Ciarán P. Kelly and J. Thomas LaMont. *Clostridium difficile* – more difficult than ever. N Engl J Med, 359(18):1932–1940, 2008.

[4] Neha Alang and Colleen R. Kelly. Weight gain after fecal microbiota transplantation. Open Forum Infect Dis, 2(1), 2015.

[5] Simone S. Li, Ana Zhu, Vladimir Benes, Paul I. Costea, Rajna Hercog, Falk Hildebrand, Jaime Huerta-Cepas, Max Nieuwdorp, Jarkko Saloj¨arvi, Anita Y. Voigt, Georg Zeller, Shinichi Sunagawa, Willem M. de Vos, and Peer Bork. Durable coexistence of donor and recipient strains after fecal microbiota transplantation. Science, 352(6285):586–589, 2016.

[6] Zain Kassam, Christine H. Lee, Yuhong Yuan, and Richard H. Hunt. Fecal microbiota transplantation for *Clostridium difficile* infection: Systematic review and meta-analysis. Am J Gastroenterol, 108(4):500–508, 2013.

[7] A. Khoruts and M. J. Sadowsky. Understanding the mechanisms of faecal microbiota transplantation. Nat Rev Gastroenterol Hepatol, 2016.

[8] Loek P. Smits, Kristien E. C. Bouter, Willem M. de Vos, Thomas J. Borody, and Max Nieuwdorp. Therapeutic potential of fecal microbiota transplantation. Gastroenterology, 145(5):946–953, 2013.

[9] Michael J. Sadowsky and Alexander Khoruts. Faecal microbiota transplantation is promising but not a panacea. Nat Microbiol, 1, 2016.

[10] Paul Moayyedi, Michael G. Surette, Peter T. Kim, Josie Libertucci, Melanie Wolfe, Catherine Onischi, David Armstrong, John K. Marshall, Zain Kassam, Walter Reinisch, and Christine H. Lee. Fecal microbiota transplantation induces remission in patients with active ulcerative colitis in a randomized controlled trial. Gastroenterology, 149(1):102–109.e6, 2015.

[11] Thomas D. Cook and David L. DeMets. Introduction to Statistical Methods for Clinical Trials. Chapman & Hall/CRC, 2008.

[12] Robert H. Bartlett, Dietrich W. Roloff, Richard G. Cornell, Alice French Andrews, Peter W. Dillon, and Joseph B. Zwischenberger. Extracorporeal circulation in neonatal respiratory failure: A prospective randomized study. Pediatrics, 76(4):479–487, 1985.

[13] Shein-Chung Chow and Mark Chang. Adaptive Methods in Clinical Trials. Chapman & Hall/CRC, 2007.

[14] Thomas Hahn. Cuba – a library for multidimensional numerical integration. Comput Phys Commun, 168(2):78–95, 2005.

[15] Ole Tange. Gnu parallel – the command-line power tool. login: The USENIX Magazine, 36(1):42–47, 2011.

[16] Donald A. Berry and Bert Fristedt. Bandit Problems: Sequential Allocation of Experiments. Springer Netherlands, 1985.

[17] James H. M. Hung, Robert T. O’Neill, Sue-Jane Wang, and John Lawrence. A regulatory view of adaptive/flexible clinical trial design. Biom J, 48(4):565–573, 2006.

[18] L. J. Wei, R. T. Smythe, D. Y. Lin, and T. S. Park. Statistical inference with datadependent treatment allocation rules. J Am Stat Assoc, 85(409):156–162, 1990.

[19] L. J. Wei and S. Durham. The randomized play-the-winner rule in medical trials. J Am Stat Assoc, 73(364):840–843, 1978.

[20] M. Zelen. Play the winner rule and the controlled clinical trial. JAm Stat Assoc, 64(325):131–146, 1969.

[21] L. J. Wei. The generalized polya’s urn design for sequential medical trials. Ann Stat, 7(2):291–296, 1979.

[22] L. Trippa, E. Q. Lee, P. Y. Wen, T. T. Batchelor, T. Cloughesy, G. Parmigiani, and B. M. Alexander. Bayesian adaptive randomized trial design for patients with recurrent glioblastoma. J Clin Oncol, 30(26):3258–3263, 2012.

[23] Sofía S. Villar, James Wason, and Jack Bowden. Response-adaptive randomization 429 for multi-arm clinical trials using the forward looking gittins index rule. Biometrics, 71(4):969–978, 2015.

[24] Noortje G. Rossen, Susana Fuentes, Mirjam J. van der Spek, Jan G. Tijssen, Jorn H.A. Hartman, Ann Duflou, Mark L¨owenberg, Gijs R. van den Brink, Elisabeth M.H. Mathus-Vliegen, Willem M. de Vos, Erwin G. Zoetendal, Geert R. D’Haens, and Cyriel Y. Ponsioen. Findings from a randomized controlled trial of fecal transplantation for patients with ulcerative colitis. Gastroenterology, 149(1):110–118, 2015.

[25] Chinyu Su, Gary R. Lichtenstein, Karen Krok, Colleen M. Brensinger, and James D. Lewis. A meta-analysis of the placebo rates of remission and response in clinical trials of active crohn’s disease. Gastroenterology, 126(5):1257–1269, 2004.

[26] Chinyu Su, James D. Lewis, Brittany Goldberg, Colleen Brensinger, and Gary R. Lichtenstein. Ameta-analysis of the placebo rates of remission and response in clinical trials of active ulcerative colitis. Gastroenterology, 132(2):516–526, 2007.

[27] C. J. Clopper and E. S. Pearson. The use of confidence or fiducial limits illustrated in the case of the binomial. Biometrika, 26(4):404–413, 1934.

[28] Y. Cheng and D. A. Berry. Optimal adaptive randomized designs for clinical trials. Biometrika, 94(3):673–689, 2007.

